# Bacterial lipids traverse the hydrophobic groove of TamB

**DOI:** 10.1101/2025.03.12.642747

**Authors:** Yiechang Lin, Ben Corry

## Abstract

The double-membrane envelope of Gram-negative bacteria protects it against environmental stress and antibiotics. Phospholipids are a core component of the bacterial outer membrane (OM). However, how phospholipids are transported to the OM from the inner membrane (IM) where they are synthesized is poorly understood. We show that the AsmA-like protein TamB transfers lipids through a large hydrophobic groove which directly bridges the inner and outer membranes. Lipid dissociation at the outer membrane is impeded when TamB is bound to its OM partner protein TamA but occurs spontaneously when TamA is not bound. Other members of the AsmA-like family in *E. coli* can also accommodate lipids within their hydrophobic grooves, suggesting they may also function as lipid transporters. Our findings highlight the lipid transport ability of TamB and other AsmA-like proteins, suggesting their importance in maintaining bacterial OM integrity.

## Introduction

Gram-negative bacteria possess a double membrane envelope which serves as a protective barrier against environmental stress and antibiotics (*1*). This envelope comprises an inner membrane (IM) and outer membrane (OM) separated by an aqueous periplasmic compartment containing a peptidoglycan layer. The IM surrounds the cytoplasm and is composed of phospholipids, predominantly phosphatidylethanolamine, phosphatidylglycerol and cardiolipin. The OM, which separates the periplasm from the external environment and plays critical roles in maintaining cell integrity, is highly asymmetric. Its outer leaflet is composed of lipopolysaccharides, which prevent the passage of hydrophobic molecules including antibiotics while the inner leaflet is composed of phospholipids.

How intracellularly synthesized phospholipids are transported to the inner leaflet of the OM has eluded our understanding until recently, when members of the AsmA-like protein family were proposed as the conduits for lipid transfer from the IM to the OM (*2–5*). In *E. coli*, this family contains 6 members (TamB, YhdP, YdbH, YhjG, YicH and AsmA), which are thought to be related to the eukaryotic repeating β-groove proteins which transfer lipids between organelles (*6*). Deletion of the genes associated with the three largest members of this family (TamB, YhdP, and YdbH) generates a lethal phenotype while deletion of *tamB* and *yhdP* together disrupts OM lipid homeostasis (*4, 5*). Furthermore, recent *in vivo* crosslinking assays and molecular dynamics simulations showed that phosphate-containing molecules directly interact with the hydrophobic groove of YhdP and phospholipids can spontaneously enter the groove during simulations (*7*). Results from these genetic, biochemical and computational studies support the idea that TamB, YhdP, and YdbH may act as redundant lipid transporters in E. coli, facilitating the spontaneous diffusion of phospholipids from the IM to OM.

The largest of these three proteins, TamB, complexes with the OM protein TamA to form the Translocation and Assembly Module (TAM) (*8*). While the β-barrel assembly machine (BAM) is responsible for catalyzing the assembly of most *E. coli* OM proteins, the structurally and evolutionarily related TAM has also been proposed to play a role in assembling several OM proteins (*8–11*). Additionally, TAM was recently shown to directly catalyze the assembly of several OM proteins in a BAM independent manner when reconstituted into liposomes (*12*). These findings raise the question of whether TAM could play dual roles in transporting lipids and assembling OM proteins, and what roles TamA and TamB each play individually during this process. Additionally, while previous studies imply a role for TAM in lipid homeostasis, it has been suggested that the phenotypes observed in these genetic studies could stem from secondary consequences of OM protein assembly (*11*), and a direct role in lipid transport has yet to be conclusively shown. Here, we use a combination of structural modelling, coarse grained and all-atom molecular dynamics simulations to investigate the lipid transfer ability of TAM and other members of the AsmA-like family. Our data suggests IM lipids can spontaneously enter and traverse the large periplasmic spanning groove of TamB in a TAM model embedded in both the bacterial IM and OM. Lipid dissociation at the OM is impeded when TamA is in complex with TamB but occurs spontaneously in structures of TamB alone. Finally, we also demonstrate that all six members of the AsmA family in *E. coli* can accommodate lipids within their hydrophobic grooves. Our results support the idea that the TAM directly transports lipids from the IM to the OM and provide insights into the mechanisms through which this occurs.

## Results

### Inner membrane lipids spontaneously traverse the hydrophobic groove of TamB

To assess the ability of the TamA/TamB complex (hereafter referred to as TAM) to act as a conduit for lipid transfer from the *E. coli* inner membrane (IM) to the outer membrane (OM), we carried out coarse grained molecular dynamics simulations of TAM. Our starting TAM model was produced by McDonnell et al. using a combination of AF2Complex and membrane morphing simulations to generate a structure that can plausibly span the bacterial envelope (*13*). In all replicate (n=5) simulations of this model with the N-terminal helix of TamB embedded in the bacterial IM (modelled as 75% POPE, 20% POPG, 5% CDL2) (*14–16*) and TamA embedded in the OM (upper: 100% ReLPS, lower: 90% POPE, 5% POPG, 5% CDL2) (*17*), we observed lipids spontaneously entering the hydrophobic groove of TamB from the upper leaflet of the IM (Fig. 1A-B, Movie S1).

**Figure 1.**
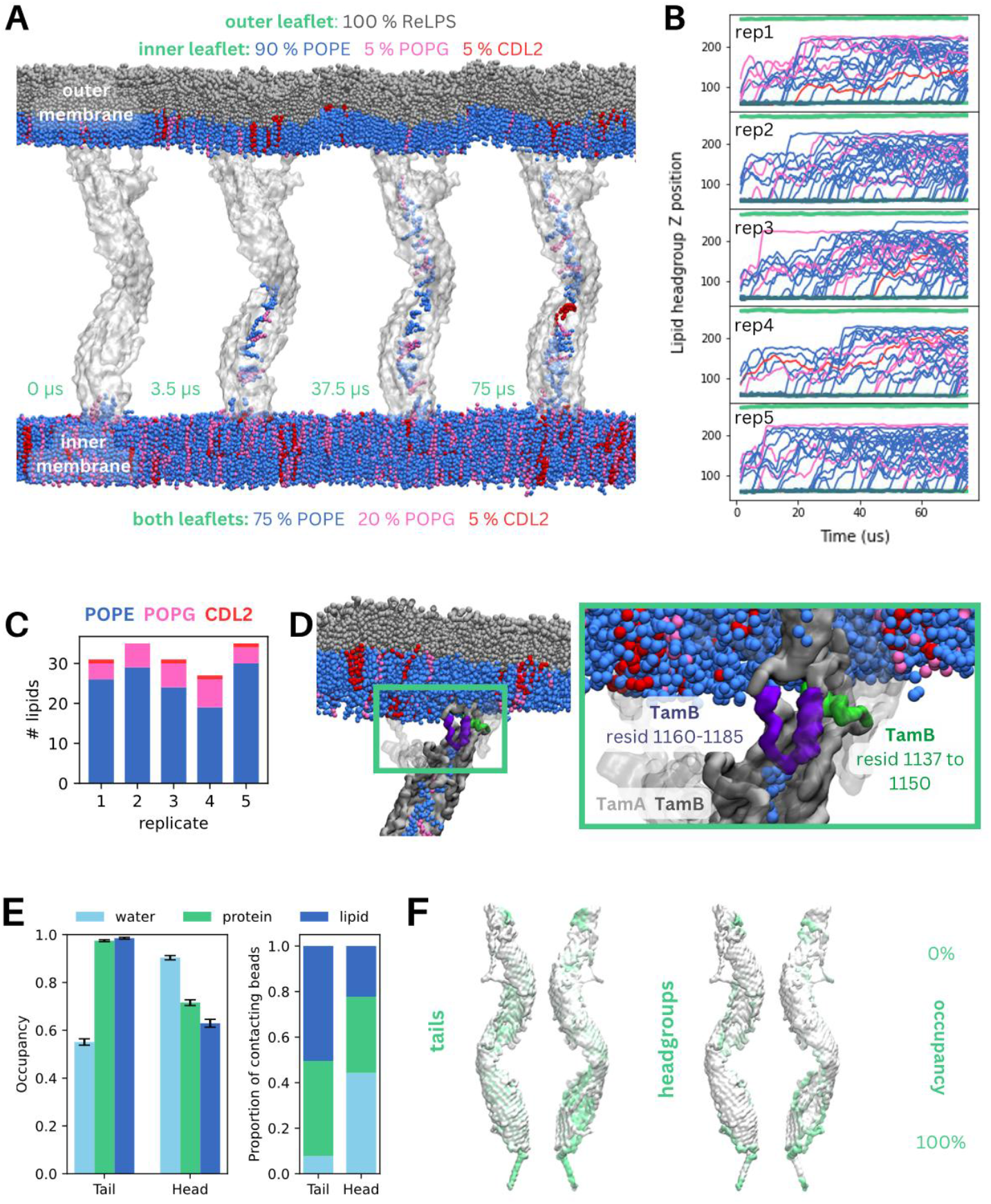
Inner membrane lipids spontaneously traverse the hydrophobic groove of TamB. **(A)** Representative snapshots of the TamA:TamB (TAM) complex embedded in the bacterial inner and outer membranes after 0, 3.5, 37.5 and 75 μs of coarse-grained molecular dynamics simulations showing lipid movement from the IM into the hydrophobic groove of TamB. Lipids are colored by type (ReLPS – grey, POPE-blue, POPG – pink, CDL2 – red). **(B)** Lipid headgroup z positions across simulation time, with each panel representing one of 5 replicates. Each line indicates the z position of one lipid headgroup, coloured by the lipid type (POPE-blue, POPG – pink, CDL2 – red). The green lines indicate the headgroup positions of lipids in the inner and outer membranes. **(C)** The number of different lipids present in the hydrophobic groove of TamB in each of five replicates after 75 μs of simulation, coloured by the lipid type (POPE-blue, POPG – pink, CDL2 – red). **(D)** Representative snapshot after 75 μs of simulation, showing a POPG lipid blocked from moving further upwards by a TamB helix (resid 1160-1185, purple), near the interface of TamA (majority of protein embedded in the OM, visible domains indicated in transparent grey) with TamB. A short helix (resid 1137-1150, green) in TamB marks the top of the hydrophobic column, which ends prior to meeting the OM. **(E)** TamB surface coloured by occupancy of lipid headgroups (left) and tails (right). Regions of higher occupancy are indicated by darker colouring.

Over the course of each 75 μs simulation, lipids filled the wide, periplasm spanning groove of TamB, traversing upwards towards the OM. By the end of the simulation, an average of 31.8±1.5 lipids were present in the hydrophobic groove of TamB (Fig. 1C). The relative proportions of lipid types within the groove closely matched the composition of the bacterial IM, with most of the lipids in the groove being POPE (Fig. 1C).

While lipids moved freely from the IM into TamB, and lipids can pass each other within the groove, we did not observe any lipid dissociation events from the groove to the OM within the timescale of our simulations. Interestingly, lipids in TamB cease to move upwards at around 10-20 Å from the OM (Fig. 1B, D). This is due to the lack of clear hydrophobic path for the lipid as the hydrophobic groove of TamB ends (Fig. 1D inset, green helix - resid 1137 to 1150) before meeting the OM. An additional bent helix (Fig. 1D inset, purple helix – resid 1160-1185) occludes upward lipid movement in this replicate, presenting another obstacle to continuous lipid flow between the IM and OM via TamB.

As lipids moved through the hydrophobic groove, their tail moieties interacted with other lipids in the groove as well as residues belonging to TamB for > 95% of simulation time (Fig.1E, left). Protein and lipid interactions formed the major proportion of contacts for lipid tails (Fig. 1E, right), with protein-tail interactions occurring on the interior of the TamB groove (Fig. 1F). In contrast, the headgroups of these lipids predominantly interacted with surrounding water and the peripheral edges and loops of the protein (Fig. 1E-F).

Spontaneous entry of inner membrane lipids into the hydrophobic groove of TamB was also observed in simulations of an unmorphed AlphaFold2 model of TamB embedded in a model IM single bilayer system (Fig S1A). This phenomenon was also observed for the morphed TamB model embedded in the IM in simulations using both the Martini 2.2 and Martini 3 forcefields (Fig. S1B-C). This suggests that the entry of lipids into TamB is independent of the structural model, the use of a single or double bilayer system, and forcefield used in our simulations.

### AF2 predictions suggests alternative orientations of TamB in the OM

Given that free lipid dissociation at the bacterial OM does not occur in our TAM simulations using a structural model where TamA and TamB interact, we wondered if alternate configurations could allow free lipid exchange at the OM. One possibility is that the conformation of TamB within the TAM complex changes such that the hydrophobic groove moves closer to the lipid bilayer. To address this, we carried out AlphaFold2 structural predictions of TAM, TamA alone, and TamB alone, generating 100 structures of each using default parameters (without alteration of the multiple sequence alignment).

In all 100 predicted TAM structures, TamA and TamB interact at the beta barrel region, with additional interactions formed between the hydrophobic groove of TamB and the first polypeptide-transport-associated (POTRA) domain in TamA (Fig. 2A). At the OM, β sheet templating between residues 266-276 of TamA and the last 10 residues (resid 1249-1259) of TamB defines the interaction interface. Compared to the crystal structure (*18*) and AF2 predictions of TamA alone (which are similar to one another, as evidenced by a TM score of 0.97), where the beta strand formed by residues 266-276 interacts instead with the final beta strand of TamA (resid 569-574), predictions of the complex show a widening of the opening in the OM beta-barrel due to interaction with TamB. All TAM structures adopted similar conformations, particularly in terms of the z-displacement of the hydrophobic groove of TamB relative to TamA (Fig. 2B). As these structures were predicted without the context of the IM and OM, the resulting structures have lengths of around 100-150 Å (distance between the center of masses of the N-terminal IM helix and the OM beta strand region), with some diversity sampled in the angle made between the IM helix and OM segments (Fig. 2C). Notably, due to the lack of structural diversity in the produced TAM structures, none showed a hydrophobic groove which could be plausibly embedded in the OM to provide a continuous pathway for lipid dissociation.

**Figure 2.**
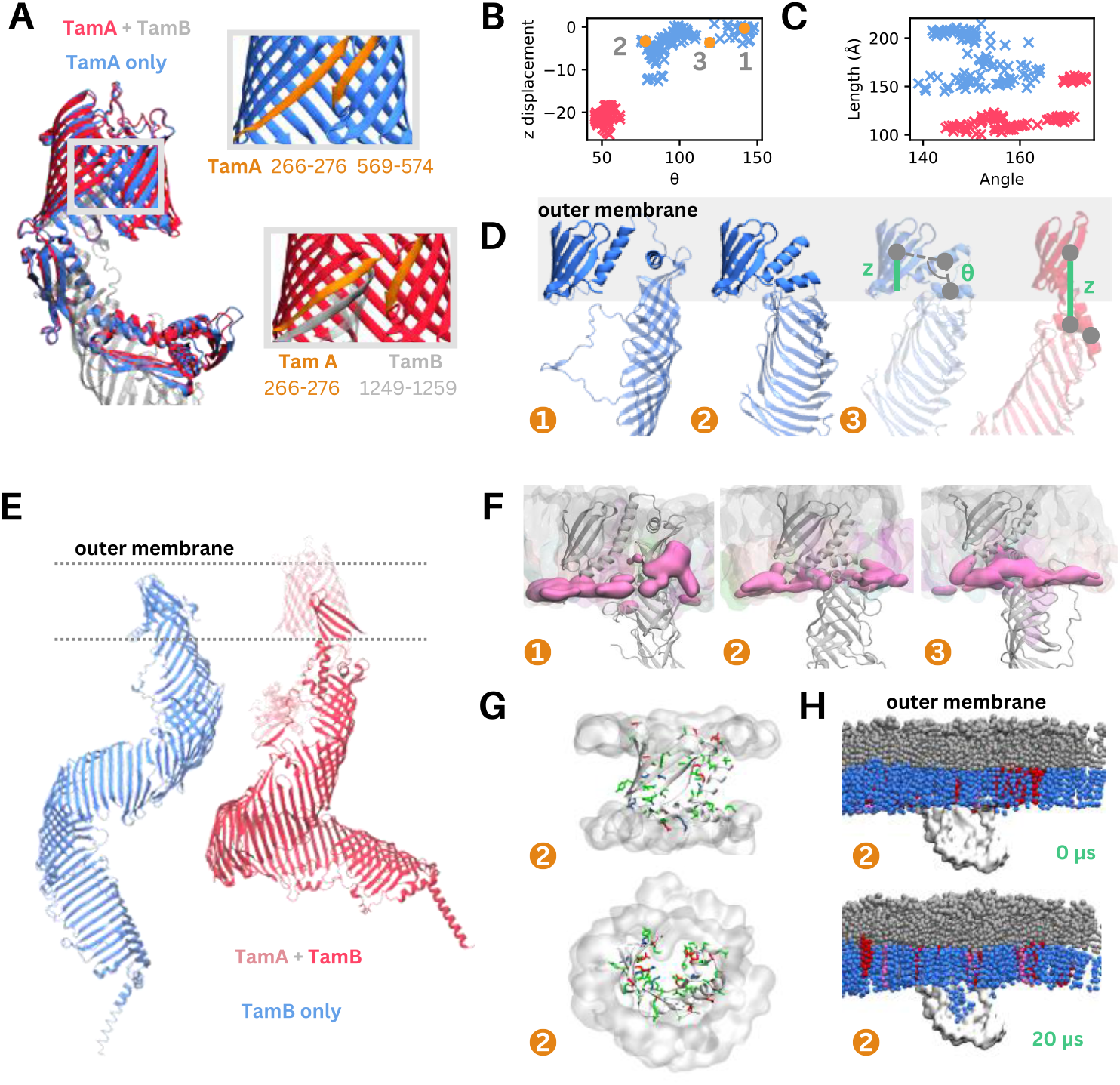
AF2 predictions suggests alternative orientations of TamB in the OM. **(A)** Overlay of the top ranked AlphaFold2 structures of TamA, predicted alone (blue) and together with TamB (red). Inset: the beta strand interface of TamA predicted alone (top), showing an interaction between two beta strands (orange, resid 272-276 and 569-574) and TamA predicted together with TamB (bottom), showing an interaction between a beta strand in TamA (orange, resid 266-276) and in TamB (grey, resid 1249-1259). **(B)** The angle made by the beta strand region (resid 1188 to 1259), OM helix (resid 1151 to 1187) and helix at the top of the hydrophobic groove (resid 1140 to 1150) plotted against the z distance between the center of masses of the beta strand region and the upper region of the hydrophobic groove. Each datapoint represents one of 100 TamB predicted alone (blue) or together with TamA (red). **(C)** The angle made between the IM helix (resid 1 to 28) and the OM beta strand region (resid 1188 to 1259) plotted against the length of TamB in each predicted structure as measured by the distance between the center of masses of the IM helix and the OM beta strand region. Each datapoint represents one of 100 TamB predicted alone (blue) or together with TamA (red). **(D)** Structural comparison of three representative structures of the OM portion of TamB predicted alone (blue) and with the OM portion of the top ranked TamA (red) with the metrics plotted in (C) indicated on the figure. The structures are numbered to correspond with the positions in (B) and reflect the structures presented in (F), (G) and (H). **(E)** The top ranked AlphaFold2 structures of TamB, predicted alone (blue) and together with TamA (red, TamA shown in transparent colouring). **(F)** Representative snapshots from all-atom simulations of the three TamB structures embedded in the bacterial OM after 500 ns of simulations. The protein is shown using cartoon representation, with the lipid membrane depicted in a transparent surface. The locations of the phosphate atoms of lipids belonging to the lower leaflet of the OM are highlighted using a pink surface. **(G)** Snapshot of the TamB beta sheet region embedded in the OM during all atom simulations viewed from the plane of the membrane (top) and from the extracellular space (bottom). Polar sidechains are shown (red: acidic, blue: basic and green: polar), and the location of the lipid headgroups near the protein is indicated by the grey surface. **(H)** Representative snapshots of TamB embedded in the bacterial OM after 0 and 3 μs of coarse-grained molecular dynamics simulations showing lipid movement from the OM into the upper segment of the TamB hydrophobic groove.

Interestingly, structures of TamB differed considerably when it was predicted alone compared to predictions in complex with TamA (Fig. 2B-E). The predicted structures of TamB alone are longer, with the longest structures having lengths of ∼215 Å, making them likely to be capable of spanning the periplasm independently (Fig. 2C). The absence of TamA in the prediction also leads to a significant change in the relative orientations of the outermost beta-sheet and alpha helix regions as well as the hydrophobic groove (Fig. 2B, D). This change in orientation (which would be sterically occluded in the presence of TamA) significantly alters which regions of TamB which would be embedded in the OM (Fig, 2D), with the structures predicted in the absence of TamA placing the hydrophobic groove closer to the OM.

Given the differences between structural predictions of TamB in the presence and absence of TamA, we wondered whether the altered orientation of the OM portion of TamB when predicted alone may represent a physiologically relevant conformation through which phospholipids can freely move between the protein and the OM. To this end, we first assessed whether the alternate conformations of TamB could stably embed in the OM, especially as there are several polar and charged residues in portions of the protein predicted to be embedded in the membrane. To do this we selected three distinct representative structures (Fig. 2C-D) and used all-atom simulations to assess the feasibility of this region to remain in the OM. After 500 ns, we observed some distortion in the lower leaflet of the OM in simulations using the top ranked (by AlphaFold2, based on average pLDDT value) structure of TamB (1) due to the high placement of the hydrophobic groove which has a polar outer surface (Fig. 2F). However, this membrane distortion was not present in simulations using structures (2) or (3) where the groove was predicted to be slightly lower relative to the OM, and no large conformational changes of the protein were observed (Fig. S2). In these stable conformations the beta sheet region of TamB spans the membrane, placing acidic residues at each end of the barrel among the lipid headgroups in each leaflet. Polar and charged residues in the middle of the sheets point inwards in a partial barrel interacting with each other or water molecules, leaving a more hydrophobic surface to associate with the lipids (Fig. 2G).

Subsequent coarse grain simulations of structure (2) showed that phospholipids belonging to the lower leaflet of the OM can spontaneously enter the upper segment of TamB. This is not the case in simulations of TAM (as discussed above) nor in simulations of TamB predicted together with TamA, with TamA removed prior to simulation (Fig. 2G). This suggests that lipid flow is not occluded by TamA directly but that when bound, it alters the conformation of TamB with respect to the OM.

Taken together, our data suggests that significantly different conformations of the TAM complex from those predicted here would be required to allow free lipid transfer to the OM in the TAM complex. In addition, our data suggests that TamB could span the periplasm without the aid of TamA and may possibly be oriented in the membrane in a manner which allows it to independently facilitate lipid flow, if correctly trafficked to and embedded in the OM.

### AsmA-like proteins may share the ability to conduct lipids

To assess how lipid transfer may differ between TamB and other members of the AsmA family in *E. coli*, we carried out 75 μs CG MD simulations of AlphaFold2 predicted structures of YhdP and YdbH, the two other members of the family suggested to transport lipids. For YhdP the predicted structure on the AlphaFold2 database contained a flexible loop near the IM which completely prevented lipid entry in our simulations, so we carried out additional simulations using an AlphaFold3 model of the protein predicted with palmitic acid in the hydrophobic groove which caused displacement of the loop. In these simulations, we observe lipids spontaneously entering the hydrophobic groove of both YhdP and YdbH from the periplasmic leaflet of the IM (Fig. 3A-B).

**Figure 3.**
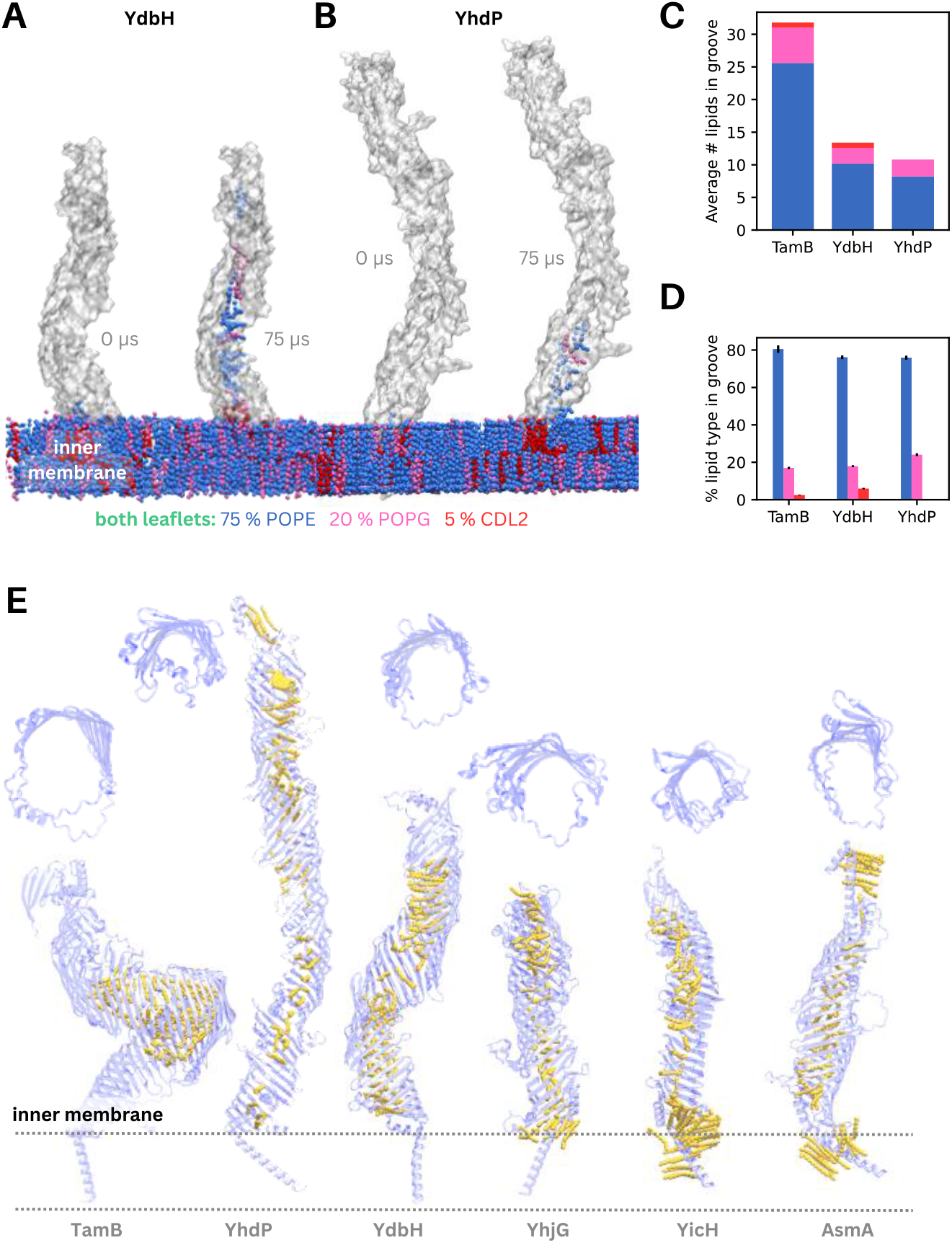
MD simulations and AF3 predictions of AsmA proteins suggest they share the ability to conduct lipids. **(A)** Representative snapshots of YdbH embedded in the bacterial inner membrane after 0 and 75 μs of coarse-grained molecular dynamics simulations showing lipid movement from the IM into the hydrophobic groove. (POPE-blue, POPG – pink, CDL2 – red). **(B)** Representative snapshots of YhdP embedded in the bacterial inner membranes after 0 and 75 μs of coarse-grained molecular dynamics simulations showing lipid movement from the IM into the hydrophobic groove. **(C)** The average number of different lipids present in the hydrophobic groove of TamB, YdbH and YhdP after 75 μs of simulation, coloured by the lipid type (POPE-blue, POPG – pink, CDL2 – red). **(D)** The % lipid composition within the hydrophobic groove of TamB, YdbH and YhdP after 75 μs of simulation, coloured by the lipid type (POPE-blue, POPG – pink, CDL2 – red). **(E)** Snapshots of TamB, YhdP, YdbH, YicH, YhjG and AsmA AF3 predictions made in the presence of 50 palmitic acid molecules (yellow). Cut though images of each protein are shown above the full structure to indicate the width of the hydrophobic groove.

The number of lipids present in the hydrophobic groove differed between TamB, YdbH and YhdP (Fig. 3C), which may be due in part to the difference in cavity volume and length between these proteins (Fig. S3A). As the protein with the widest hydrophobic groove and largest cavity volume (Fig. S3), TamB was able to accommodate more than double the number of lipids when compared to YdbH and YhdP. During the 75 μs long simulations, lipids fully traversed the groove of YdbH and reached the top of the protein. In contrast, owing to the narrow constriction in the YhdP groove (Fig. S3B), lipids were prevented from fully occupying the length of YhdP. We note that the shorter length of YdbH, which does not span the periplasm alone, and the presence of the constriction in YhdP (Fig. S3B), partially accounts for the reduced number of lipids within the groove when compared to TamB.

The proportion of each lipid type found within the grooves of TamB, YdbH and YhdP mostly matched the composition of the IM (Fig. 3D), although few cardiolipin molecules were found in the grooves of TamB and YdbH, and none entered YhdP in any replicate.

Finally, given that AlphaFold3 was able to correctly place palmitic acid within the hydrophobic groove of YhdP, we carried out predictions for all 6 AsmA proteins with 50 palmitic acid molecules and showed that these are placed within the grooves of each protein (Fig. 3E). Taken together, these data suggest that the hydrophobic groove of all 6 proteins can accommodate lipids.

## Discussion

The AsmA-like family of proteins were recently proposed to act as conduits for lipid transfer between the bacterial IM and OM. One member of this family, TamB, interacts with an OM protein TamA to form the Translocation and Assembly Module (TAM). TAM is known to aid in the assembly of several OM proteins (*8–12*), in addition to being implicated in lipid transport (*4, 5*). We sought to directly investigate whether TAM transports lipids and elucidate the individual roles of TamA and TamB in this process. In simulations, phospholipids spontaneously enter and traverse the large hydrophobic groove of TamB, strongly suggesting its role in lipid transport. Additionally, we show that TamB alone may be able to facilitate lipid transfer if it can be trafficked to the OM and inserted such that the hydrophobic groove is continuous with the lower leaflet of the OM, although we cannot exclude the possibility that other proteins, including TamA, could play roles in offloading lipids at the OM. Finally, we show that all 6 members of the AsmA-like family can accommodate lipids in their hydrophobic grooves.

The TAM structures predicted using AF2 in this study and by others always place the complexation interface such that the OM segment of TamB completes the TamA β-barrel. This is consistent with previous size-exclusion chromatography experiments which show that residues 22-293 of TamA interact with residues 1180-1259 of TamB, with the final β-strand of TamB (residues 1252-1259) being important for this complex formation (*8, 13*). This suggests TamA and TamB likely bind to each other at the OM mediated by interactions between β-strands belonging to both proteins, and the TAM structure simulated here likely represents one plausible configuration of the two proteins in bacterial cells. Our use of a morphed AlphaFold2 model was necessary to enable correct placement of the TAM complex, consistent with the proposed orientation of TamA in the OM and TamB in the IM. While this awaits validation from cryo-EM structures of the full TAM complex, which could also help to clarify the native interaction interface between TamA and TamB, additional simulations using an unmorphed AF2 model shows that lipid entry into the hydrophobic groove occurs irrespective of the model used, suggesting this is not an artefact of the model.

Interestingly, while lipid influx from the IM and traversal through the hydrophobic groove of TAM was rapid, lipids did not dissociate at the OM within the timescale of our simulations. In contrast, structural predictions of TamB alone suggest that it may be able to span the periplasm on its own, and simulations of one of these structures in the OM showed that these conformations were stable and allowed spontaneous movement of phospholipids from the OM into the groove. These results suggest that it may be possible for TamB to facilitate phospholipid movement from the IM to the OM without TamA bound during this process. However, previous genetic knock out experiments have shown that TamA is required for TamB’s roles in OM lipid homeostasis, with the same phenotypes being observed when tamA or tamB is knocked out in bacteria lacking other lipid transporters (*4, 19*).

There are several explanations which can account for these experimental and simulation data. Firstly, it’s possible that lipid dissociation from TamB at the OM can occur while TamB is in complex with TamA in an alternative conformation not captured by our AlphaFold modelling or at timescales beyond what we can currently simulate. Secondly, it is also plausible that one or more additional accessory proteins aid in lipid offloading at the OM. Alternatively, our observations may be explained by a model in which TamA aids correct localization and folding of TamB at the OM, but where the two proteins are not associated during the lipid transfer process. This proposed transient disassociation of TamA from TamB is perhaps supported by recent work which suggests that TamB dissociates from the TamA β-barrel during OMP assembly as the interaction interface is the site for β -strand templating of other proteins (*12*).

Comparison of our simulations of TAM, YhdP and YdbH reveal differences in the number of lipids which can be accommodated in the hydrophobic grooves of these proteins, with the wider groove of TamB permitting more than twice as many lipids than YhdP and YdbH to enter. These findings are consistent with the proposal that TamB may facilitate the trafficking of nascent OMPs to the OM via this hydrophobic groove, perhaps simultaneously allowing lipid transport (*12, 20*). YhdP is capable of spanning the periplasm alone and is longer than YdbH, which requires the aid of additional proteins to completely bridge the space between the IM and OM (*7, 21*). Despite this, fewer lipids were present in the groove of YhdP at the end of our simulations, owing to the narrow nature of the groove and the presence of a constriction in the groove in the middle of the protein in our predicted structure.

Our simulations of YhdP are consistent with previous findings that phosphate-containing molecules (ostensibly phospholipids) span the length of the YhdP groove, supporting its proposed role in lipid transport (*3–5, 7*). CG simulations spanning 5 µs captured on average one lipid entering the hydrophobic groove from the IM, fewer than the 3-10 lipids were observed in the same timespan owing to the use of a different starting structure with the N-terminal loop displaced (*7*). Interestingly, spontaneous bilayer formation simulations conducted with the C-terminal segment of YhdP showed that the protein interacted only the inner leaflet of the OM and no lipid entry events were observed from the OM (*7*). This finding might support the idea that an additional accessory protein could be required to offload lipids at the OM across the different transporters. Alternatively, the YhdP conformation used, or the timescale of the simulations could also account for this result.

Previous genetic studies showed that only one of TamB, YhdP or YdbH is required for bacterial survival (*4, 19*), and the purpose of multiple redundant lipid transporters remains an open question. The possibility of differential phospholipid-substrate transport preferences between YhdP, TamB and YdbH has been proposed to explain the existence of three redundant lipid transporters and differences in phenotypes when these proteins were knocked out (*22*). These differences were based more strongly on the saturation of the fatty acyl chains rather than lipid headgroups, with YhdP proposed to transport cardiolipin and saturated lipids while TamB was proposed to transport more unsaturated lipids. While the data from our simulations are insufficient to draw strong conclusions on this point, there does not appear to be any obvious selectivity for or against a specific lipid headgroup type by any of these proteins, nor a large difference between the types of lipids found in the hydrophobic groove when comparing between proteins, although we note that cardiolipin was not found in the hydrophobic groove of YhdP in any of our simulations. Further investigation of TamB, YhdP and YdbH is necessary to determine whether these proteins exhibit differences in substrate specificity or could play distinct roles physiologcially. In addition, much remains to be understood about how AsmA-like proteins work alongside the Pqi, Let and Mla systems which are also implicated in bacterial phospholipid transport (reviewed in (*23*)). Further study is required to better understand the directionality of lipid transport, whether energy input is required, substrate specificity and the broader physiological roles of these pathways.

Our findings offer new insights into lipid transport mechanisms in bacteria and the role of TamB in outer membrane biogenesis. Given the essential nature of the AsmA-like proteins in bacterial survival, increasing our understanding of how they function is foundational to the development of antibiotics targeting these proteins.

## Materials and Methods

### Coarse grained (CG) molecular dynamics simulations

All CG simulations were prepared with the CHARMM-GUI Martini Bilayer Builder with the Martini 2.2 forcefield with elastic networks imposed (elnedyn22) and carried out using GROMACS 2023 (*24*).

CG simulations of full-length TAM, YdbH and YhdP embedded in bacterial membranes were carried out to investigate their lipid transfer ability.

Simulations of the TAM complex were initiated from a structure produced by McDonnell et al. (2023) using a combination of AlphaFold2 predictions and membrane morphing simulations (*13*). Two separate systems containing the TAM complex were generated and manually merged. The first system contained the N-terminal helix of TamB embedded in a 18 x 18 nm bilayer representing the bacterial IM with 75% POPE, 20% POPG and 5% CDL2 in both leaflets. The total number of lipids in the IM is 1040, 520 in each leaflet. The second system contained TamA embedded in a 18 x 18 nm bilayer (700 lipids total) containing 100% ReLPS in the outer leaflet and 90% POPE, 5% POPG and 5% CDL in the inner leaflet to represent the OM. The total number of lipids in the OM is 700, with 200 ReLPS molecules (which have a larger area per lipid given the 4 tails in Lipid A) and 500 lipids in the lower leaflet. The two systems were merged by aligning the protein complex and adding the inner membrane lipids to the second system and removing the overlapping water molecules in the second system. A timestep of 15 fs was used in production simulations and 5 replicates were carried out for 75 µs each.

To investigate whether lipid entry and traversal of the TamB groove can occur using an alternative structural model, we set up systems containing an unmorphed AlphaFold2 model of TamB embedded in a model IM as described above and simulated these systems in triplicate for 18 µs each.

In addition, to ensure that lipid entry and traversal were not an artefact of the double bilayer system and/or forcefield used, we carried out simulations with just TamB from the membrane morphed structure using the Martini 2.2 and Martini 3 forcefields. As Martini 3 CDL2 parameters have yet to be implemented in the CHARMM-GUI bilayer builder, we used an 18 x 18 nm membrane composed of 80% POPE and 20% POPG in these simulations, which were also carried out in triplicate for 18 µs each.

The predicted structure of *E. coli* YdbH (UNIPROT ID: P52645) was obtained from the AlphaFold Protein Structure Database (*25, 26*). The N-terminal helix was embedded into a symmetric bilayer mimicking the lipid composition of the bacterial IM as described above. A timestep of 20 fs was used in production simulations and 5 replicates were carried out for 75 µs each.

The predicted structure of E. coli YhdP (UNIPROT ID: P46474) obtained from the AlphaFold Protein Structure Database contains an N-terminal loop which prevents unencumbered lipid access from the periplasmic leaflet of the IM. Prediction of the YhdP structure using AlphaFold2 and AlphaFold3 revealed that this loop is flexible (with multiple conformations present between predictions) and could be displaced from a blocking conformation by predicting YhdP together with palmitic acid molecules. For our simulations of YhdP, we thus used an AF3 predicted structure of YhdP in which the N-terminal loop was displaced using palmitic acid molecules. These molecules were removed prior to simulation and the N-terminal helix embedded into a bilayer mimicking the composition of a bacterial IM. A timestep of 20 fs was used in production simulations and 5 replicates were carried out for 75 µs each.

Simulations of the C-terminal portion of TamB were carried out to investigate whether phospholipids spontaneously enter from the OM. These simulations were initiated from a AF2 structural prediction of TamB (truncated, residues 900-1259). A timestep of 20 fs was used in production simulations and 3 replicates were carried out for 3 µs each.

Each system was solvated, neutralized and ionized using 150 mM NaCl. Following the standard CHARMM-GUI protocol, the system was subjected to a short energy minimization, followed by a 5-step equilibration protocol spanning 4.75 ns in total in which restraints on the protein (1000 kJ mol^-1^ nm^-2^, 500 kJ mol^-1^ nm^-2^, 250 kJ mol^-1^ nm^-2^, 100 kJ mol^-1^ nm^-2^, 50 kJ mol^-1^ nm^-2^) and lipid headgroups (200 kJ mol^-1^ nm^-2^, 100 kJ mol^-1^ nm^-2^, 50 kJ mol^-1^ nm^-2^, 20 kJ mol^-1^ nm^-2^, 10 kJ mol^-1^ nm^-2^) were gradually reduced. Production simulations were carried out in the NPT ensemble, with a pressure of 1 bar maintained using a Parinello-Rahman barostat with semi-isotropic conditions (*27*). The temperature was maintained at 310 K using a v-rescale thermostat (*28*). To prevent unrealistic protein deformation in the absence of an OM or OM anchoring region, we imposed weak backbone position restraints of 5 kJ mol^-1^ nm^-2^ in production simulations of all single bilayer systems.

### Structural Predictions

To investigate the complexation interface between TamA and TamB and in search of a TamB structure which may plausibly span the periplasm alone, E. coli TamA, TamB and TamA+TamB together (TAM) were predicted using Colabfold (*25, 29*). 100 structures (5 structures x 20 random seeds) were predicted in each case without relaxation or modification to the multiple sequence alignment, using 3 recycles.

To investigate whether the hydrophobic grooves of AsmA-like proteins are likely to accommodate lipids, the structures of *E. coli* TamB (UNIPROT ID: P39321), YhdP (UNIPROT ID: P46474), YdbH (UNIPROT ID: P52645), YhjG (UNIPROT ID: P37645), YicH (UNIPROT ID: P31433) and AsmA (UNIPROT ID: P28249) were each predicted with 50 molecules of palmitic acid using the AlphaFold3 online server (*30*).

### All Atom Simulations

All atom molecular dynamics simulations were carried out using GROMACS 2023 with the CHARMM36 forcefield (*24, 31*). Simulation systems prepared using the CHARMM-GUI Bilayer Builder with three representative starting structures of TamB generated by AlphaFold2. These structures were truncated to include only the C-terminal portion (residues 900-1259) and embedded in a membrane containing PVCL2, PMPE, PMPG, PVPE and PVPG in a ratio of 2:8:1:8:2 in the lower leaflet and lipid A in the upper leaflet according to the protocol in Li *et al*. (2022) (*32*). The system was solvated, neutralized and ionized with 150 mM NaCl. Following the standard CHARMM-GUI protocol, the system was subjected to a short energy minimization, followed by a 5-step equilibration protocol spanning 11.25 ns in total in which restraints on the protein backbone (2000 kJ mol^-1^ nm^-2^, 1000 kJ mol^-1^ nm^-2^, 500 kJ mol^-1^ nm^-2^, 500 kJ mol^-1^ nm^-2^, 10 kJ mol^-1^ nm^-2^), sidechain (2000 kJ mol^-1^ nm^-2^, 1000 kJ mol^-1^ nm^-2^, 500 kJ mol^-1^ nm^-2^, 0 kJ mol^-1^ nm^-2^, 0 kJ mol^-1^ nm^-2^) and lipid headgroups (1000 kJ mol^-1^ nm^-2^, 500 kJ mol^-1^ nm^-2^, 500 kJ mol^-1^ nm^-2^, 500 kJ mol^-1^ nm^-2^, 0 kJ mol^-1^ nm^-2^) were gradually reduced. Periodic boundary conditions were applied in all directions. The temperature was maintained at 310 K using the Nose-Hoover thermostat (*33*). Production simulations used a timestep of 2 fs. Hydrogen bonds were constrained using the LINCS algorithm (*34*). Electrostatic interactions were treated using the Particle Mesh Ewald algorithm with a cut off of 12 Å (*35*). A van der Waals radius of 12 Å was used. Simulations were carried out in triplicate for 500 ns each.

### Analysis and Visualization

Analysis scripts were written in python using the MDAnalysis, pandas and scipy libraries (*36–38*). All simulations were visualized and snapshots prepared using Visual Molecular Dynamics (VMD) (*39*).

All analysis was performed on simulation trajectories post-processed to account for periodic boundary conditions, with the protein aligned to the first frame of the simulation. The trajectory was strided such that each frame represents 75 ns of simulation time for TamB simulations and 100 ns all other simulations. The z coordinates of lipid headgroups (PO4 beads) were tracked across the simulation time. The number of lipids present in the hydrophobic grooves of each protein was calculated as the number of headgroup (PO4, PO41) beads with z coordinates above the average z coordinate the upper leaflet headgroups of the IM. Lipid headgroup and tail binding occupancies were calculated per residue by computing the proportion of simulation time for which any bead belonging to a phospholipid headgroup (PO4, GL0, GL1, GL2, NH3, PO41, GL11, GL21, PO42, GL12, GL22) or tail moiety (all other beads) could be found within 7 Å of the residue.

To calculate the cavity volume of YhdP, YdbH and the morphed TamB model, we used the MOLEonline server to model the pathway through each protein, using ‘pore mode’ with a probe radius of 45 and interior radius of 0.8 (parameters previously used to investigate YhdP) (*7, 40*). We estimated the volume of each tunnel, which was represented by overlapping spheres, using a Monte Carlo integration method. Sphere centers and radii were extracted from the VMD script provided by the MOLEonline server. A bounding box encompassing all spheres was computed, and 10,000,000 random points were sampled within it. The tunnel volume was estimated as the bounding box volume multiplied by the fraction of points found inside at least one sphere. The length was calculated as the distance between the ends of the first and last sphere forming the tunnel. Pore radius profiles were generated by projection of the sphere centers to the tunnel axis, and plotting the corresponding radii as a function of position along the axis.

## Acknowledgments

The authors thank Dr. Denisse Leyton and Dr. Matthew Doyle for helpful discussions. This research was undertaken with the assistance of resources and services from the National Computational Infrastructure (NCI), which is supported by the Australian Government.

