## Supporting Information for "Bacterial lipids traverse the hydrophobic groove of TamB"

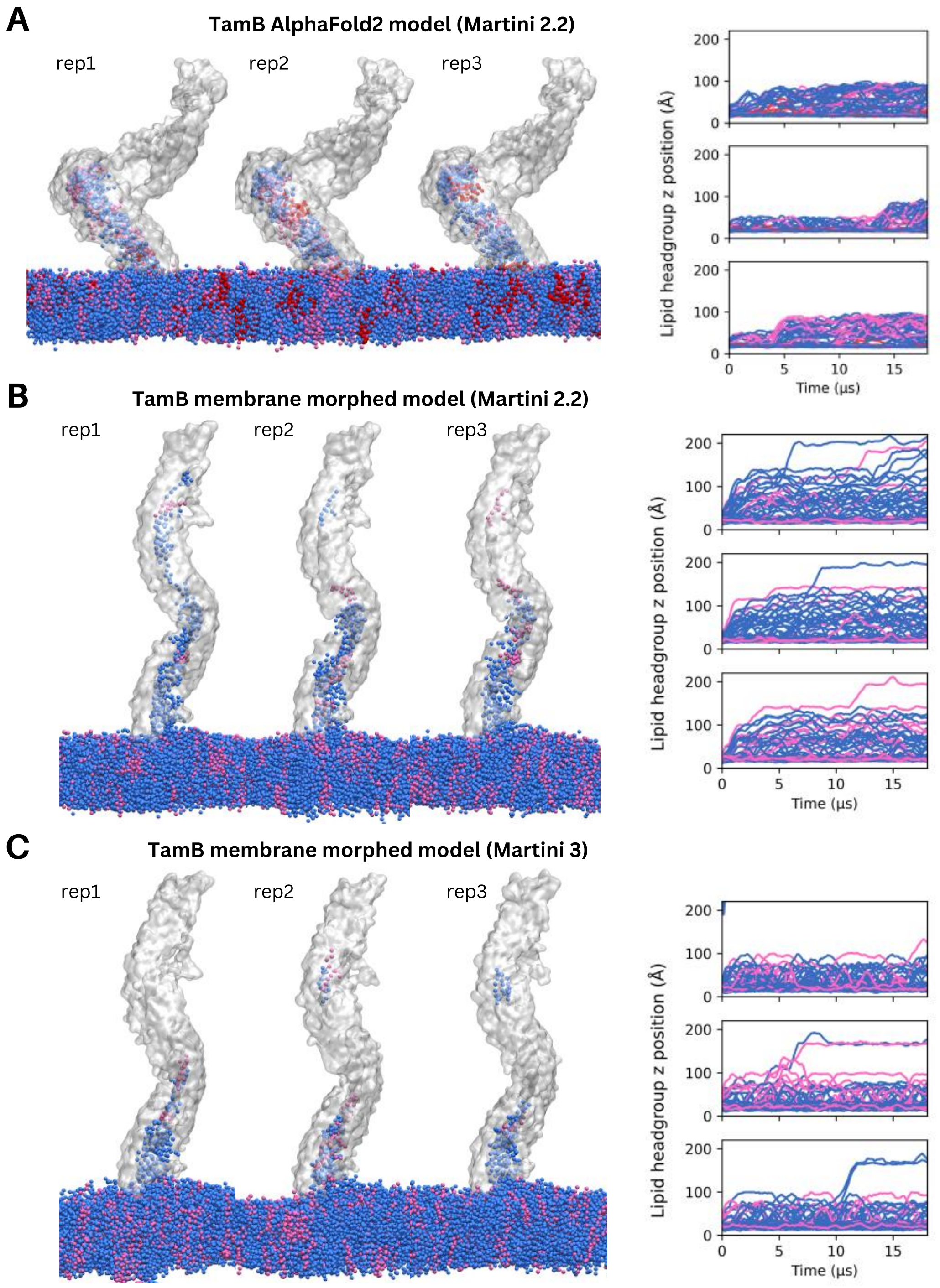


**Fig. S1. Lipid entry into TamB in single bilayer systems**

- **(A)** Final frame snapshots (left) and lipid headgroup z position plots (right) for triplicate 18 µs simulations of the AlphaFold2 model of TamB, carried out using the Martini 2.2 forcefield.
- **(B)** Final frame snapshots (left) and lipid headgroup z position plots (right) for triplicate 18 µs simulations of the membrane morphed model of TamB, carried out using the Martini 2.2 forcefield.
- **(C)** Final frame snapshots (left) and lipid headgroup z position plots (right) for triplicate 18 µs simulations of the membrane morphed model of TamB, carried out using the Martini 3 forcefield.

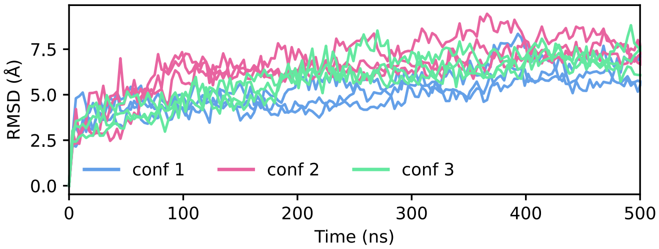


**Fig. S2.** Root mean square deviation (RMSD) plots for three conformations of the TamB OM segment in triplicate 500 ns all-atom simulation.

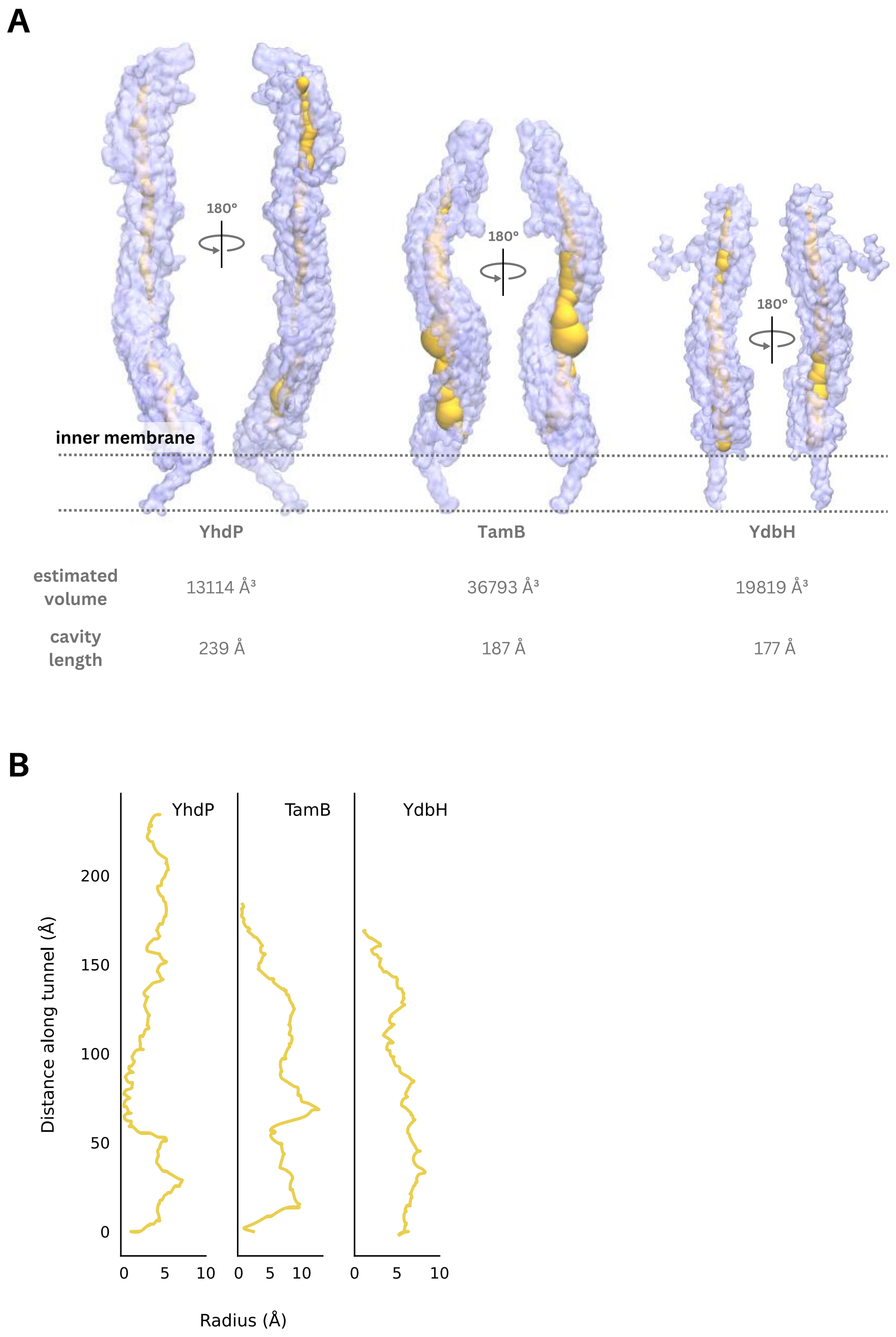


**Fig. S3. Cavity analysis of YhdP, TamB and YdbH**

- **(A)** Visualisation of the internal cavities of YhdP, TamB and YdbH. Protein surface shown in transparent purple and tunnel through the cavity (yellow) as calculated by the MOLEonline server. The estimated volume and length of each tunnel is provided.
- **(B)** 2D tunnel radius profile for each of YhdP, TamB and YdbH

**Movie S1 (separate file).** A 75 μs coarse grained simulation of TAM embedded in the E. coli inner membrane (IM) and outer membrane (OM) showing lipid movement from the IM into the hydrophobic groove of TamB. Lipids are colored by type (ReLPS – grey, POPE- blue, POPG – pink, CDL2 – red).
